# MLIB: an easy-to-use Matlab toolbox for the analysis of extracellular spike data

**DOI:** 10.1101/2025.03.25.645246

**Authors:** Maik Christopher Stüttgen

## Abstract

The analysis of neurophysiological data obtained from extracellular recordings is usually performed using a number of standard techniques. These include a) the extraction of action potentials from voltage traces and their subsequent classification, i.e., spike sorting, b) the visualization of activity, e.g., by constructing raster plots, peri-stimulus time histograms (PSTHs), and spike density functions, and c) the quantification of neuronal responses according to experimental variables such as stimulation or movement. Here I present a Matlab toolbox containing functions for the visualization and analysis of neuronal spike data. The toolbox consists entirely of one-liners that operate on vector or matrix inputs, i.e., spike and event timestamps or waveform samples. The toolbox functions provide both basic (constructing PSTHs, computing waveform characteristics etc.) and more advanced functionality, such as dimensionality reduction of multi-neuron recordings.

While offering a high degree of versatility, the toolbox should also be accessible to newcomers to neurophysiology, such as (under)graduate students or PhD students. The functions are streamlined, easy to use, and each function is extensively introduced with several examples using real or simulated data. In addition, many functions provide fully formatted plots on request, even with minimal Matlab knowledge.

## Introduction

The extracellular observation of neurons’ electrical activity constitutes one of the most powerful methods to investigate brain function. One hundred years after the first recordings of extracellular action potentials in mammals (Adrian, 1926), researchers can now record up to several thousand neurons simultaneously (Kleinfeld et al., 2019; Steinmetz et al., 2021), providing an unprecedented opportunity to investigate the workings of the brain in real time. Insights gained from this technique include the principles of neural information processing such as rate and temporal coding, feature selectivity, adaptation, and gain modulation (Adrian, 1926; DeCharms & Merzenich, 1996; Lettvin et al., 1959; Salinas & Thier, 2000). In consequence, today extracellular spike recordings are a cornerstone of neuroscience research, and are applied to a wide variety of research questions, including sensory processing, attention and cognition, decision-making, sensorimotor control, learning and memory, as well as applied research directed towards the development of brain-computer interfaces and neuroprosthetics.

The analysis of neurophysiological data obtained through extracellular recordings is typically conducted using a range of standard techniques. These include a) spike extraction and sorting, i.e., the close inspection of spike waveforms to identify and eliminate artifacts, and the waveforms’ subsequent clustering to obtain single-unit spike trains from multi-unit recordings; b) the visualization of neural activity, for example through the construction of raster plots, peri-stimulus time histograms (PSTHs), and spike density functions (SDFs); and c) the quantification of neural responses according to experimental variables such as stimulation or movement to test research hypotheses. Such analyses are usually performed with custom-written code, predominantly written in Matlab or Python, although many labs also rely on either commercially available software packages or freely available toolboxes (reviewed in (Unakafova & Gail, 2019)).

In many labs, neuronal spike data is obtained frequently by master or PhD students with little or no programming experience and limited knowledge of typical analysis procedures for neuronal spike data. (In my experience, even many experienced postdoctoral research fellows struggle with the analysis of spike data.) To provide such users with an accessible, easy-to-use solution for most standard analysis scenarios, I created a Matlab toolbox specifically for that purpose. The toolbox contains 17 functions to visualize and analyze neuronal spike data, and three additional functions pertaining to local field potential data. The toolbox functions not only cover the most basic spike analyses, such as constructing raster plots, PSTHs, and SDFs, but also more advanced and specialized analyses, among them time-resolved multiple regression of firing rates on experimental variables, clustering of multiple neurons’ PSTHs to categorize neurons into subclasses, and dimensionality reduction of multi-neuron recordings to visualize population activity. By design, the functions are easy to use for newcomers: all functions can be called directly from Matlab’s Command Window, and the use of each function is illustrated with several examples, drawing on real data recorded in rats, mice, and pigeons.

## Materials and Methods

The MLIB functions have been developed over the past decade and tested using Matlab R2023b on a standard desktop PC running Windows 11. The example data are in part taken from previous studies and include spike recordings performed in rats, mice, and pigeons (Kasties et al., 2016; Lengersdorf et al., 2014; Starosta et al., 2014; Stoilova et al., 2020; van der Bourg et al., 2017; Vandevelde et al., 2023; Yeganeh et al., 2022). Table 1 lists all MLIB functions in alphabetical order and provides brief descriptions. Beyond the basic Matlab installation, some functions require the Statistics and Machine Learning Toolbox; one function (mtune.m) requires the Curve Fitting and Signal Processing Toolboxes. An extended description of the data files, their variables, and the functions can be found in a documentation file (see Section “Data, scripts, code, and supplementary information availability” below). All experiments performed on animals were conducted in accordance with the German guidelines for the care and use of animals in science as well as with the Directive 2010/63/EU of the European Parliament and of the Council of 22 September 2010 concerning the care and use of animals for experimental purposes, and all procedures were approved by national ethics committees of the states of North Rhine-Westphalia (LANUV) and Rhineland-Palatinate (LUA), Germany (Az. 8.87-50.10.37.09.277, G 19-1-094, G 19-1-085).

**Table 1:** Overview over MLIB functions in alphabetical order.

| Function name | Primary input data | Brief description |
| --- | --- | --- |
| **mcc**<sup>1</sup> | spike waveforms from two or more clusters; spike time stamps | visualizes unit clusters and provides quality metrics to quantify spike sorting success |
| **mcheck**<sup>1</sup> | spike waveforms from one unit cluster; spike time stamps | visualizes one unit's waveforms and provides quality metrics |
| **mcluster**<sup>1</sup> | PSTHs from different units | clusters PSTHs from different units according to their similarity |
| **mcsd** | LFP data from a linear electrode array, averaged over multiple identical trials | performs a current source density analysis and plots the result |
| **mcvar** | spike time stamps from a single unit | computes the coefficient of variation for interspike intervals and the Fano factor for spike counts |
| **mdimreduct**<sup>1</sup> | PSTHs from different units | performs dimensionality reduction of spike responses from several units recorded under the same conditions |
| **mep**<sup>1</sup> | continuous local LFP data from a single or multiple electrodes | returns the evoked potential with error bars and confidence intervals |
| **mlat**<sup>1</sup> | spike and event time stamps | provides several measures of response latency |
| **mmi**<sup>1</sup> | spike counts obtained from the same unit under two different conditions | computes mutual information of spike counts |
| **mmi_psych**<sup>1</sup> | behavioral performance indices (hit rate and false alarm rate) | expresses behavioral performance in bits to compare with neuronal MI |
| **mmultregress**<sup>1</sup> | spike and event time stamps | performs a sliding-window multiple regression analysis of spike responses |
| **mnspx** | spike and event time stamps | provides a vector of spike counts in an interval around a series of events |
| **mpsth** | spike and event time stamps | constructs a PSTH and raster plot |
| **mroc**<sup>1</sup> | spike counts obtained from the same unit under two different conditions | computes the area under the ROC curve (AUROC) for two spike count distributions |
| **msdf**<sup>1</sup> | PSTH from one unit | constructs a spike-density function from a PSTH |
| **mspectan** | continuous LFP data | performs spectral analysis of LFP data |
| **mtune**<sup>2,3</sup> | spike and event time stamps | constructs and fits a tuning curve |
| **mvecstrength** | spike and event time stamps | computes vector strength of a spike response relative to a periodic signal |
| **mwa**<sup>1</sup> | spike and event time stamps | computes a 'window analysis', i.e., AUROC of a spike train in moving windows relative to a fixed reference event |
| **mwa2**<sup>1</sup> | spike and event time stamps | computes a 'window analysis', i.e., AUROC of two spike trains e.g., relative to two different stimuli |
| **mwave** | spike waveforms from a single unit | plots the average spike waveform and computes its characteristics (peak-to-trough duration etc.) |

1 requires the Statistics and Machine Learning Toolbox
2 requires the Curve Fitting Toolbox
3 requires the Signal Processing Toolbox

MLIB is designed to allow conducting basic electrophysiological analyses with a minimum of MATLAB knowledge. The library does not require installation, the user simply needs to copy the functions and example data files into a folder and then add that folder to the MATLAB user path. All functions are one-liners and operate directly on vector or array input. Customization of the functions’ outputs is achieved through the provision of input arguments, which are explained in each function’s introductory comments (type ‘doc’ and then the name of the function to read it). Additionally, a function reference script with several usage demonstrations for each function is provided to facilitate quick understanding of how the functions work. All results figures in this manuscript can be reproduced using the reference script.

For a broad introduction into the analysis of neuronal signals in general, the reader is referred to a freely available Matlab tutorial (Chakrabarti, 2024).

## Results

The analysis of extracellular spike recordings typically involves a series of analysis steps which are outlined in Figure 1. I will introduce several MLIB functions with reference to the analysis flow shown in this figure. Note that each function has features not illustrated in this paper; the full scope is described in detail in each function’s documentation and illustrated with example data in an accompanying function reference script (MLIB_Function_Reference.m).

**Figure 1.**
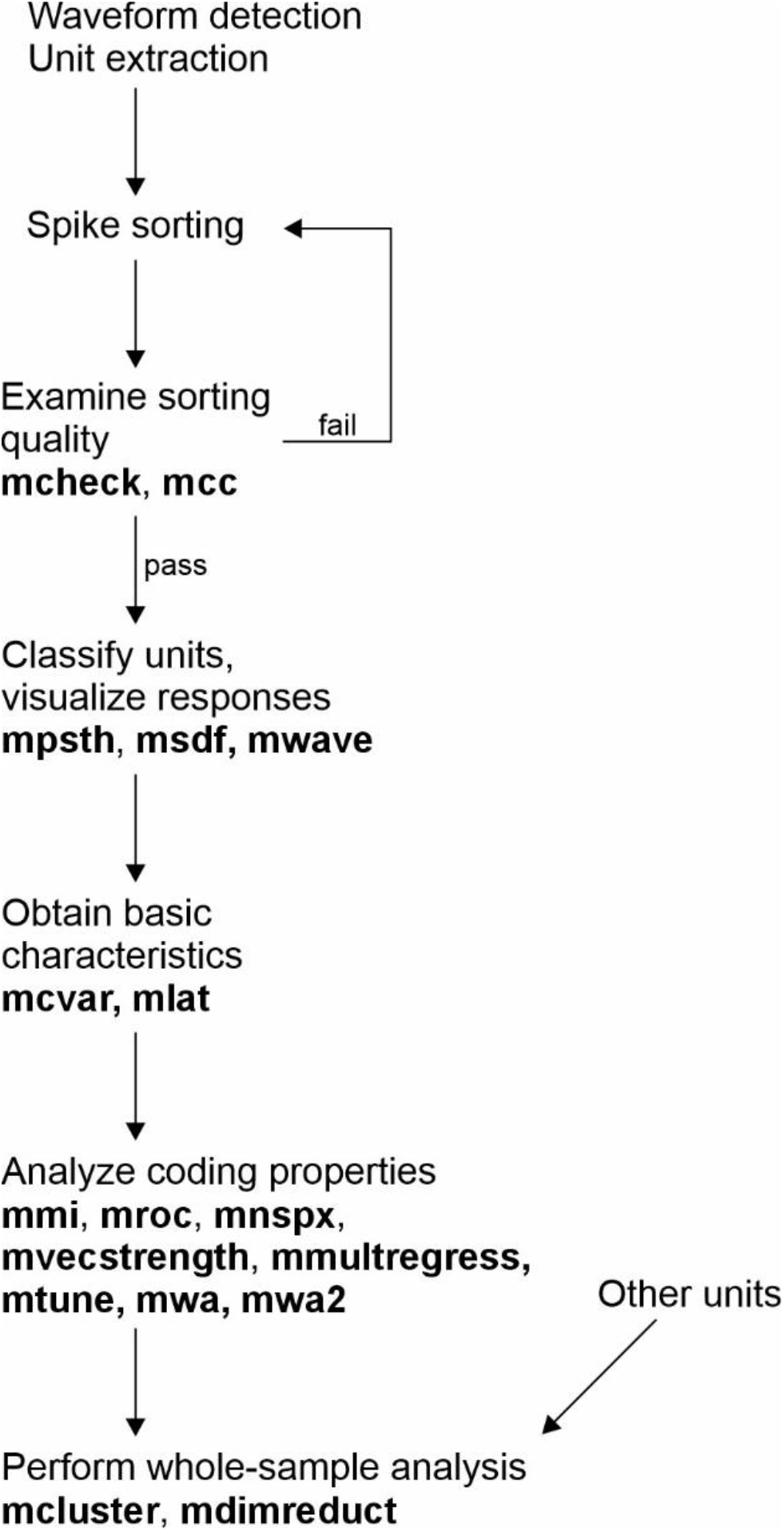
Flow chart for single-unit analysis and which MLIB functions to use at what stage of analysis.

### Checking the quality of spike sorting

The MLIB toolbox does not contain functions for spike sorting per se, as there are already multiple excellent free spike sorting packages available (Buccino et al., 2020; Hill et al., 2011; Pachitariu et al., 2024; Quian Quiroga et al., 2009; Rossant et al., 2016), in addition to some commercial software packages (e.g., Plexon’s Offline Sorter and CED’s Spike2). That said, MLIB provides functions which aid in assessing the quality of sorting post-hoc by providing several useful analyses.

These functions are illustrated in Figure 2 which shows several panels taken from the MLIB functions mcheck, mcc, and mwave. Sample waveforms of an example single unit recorded in the pigeon nidopallium caudolaterale (NCL; (Starosta et al., 2014)) are shown in Figure 2A, along with the mean and ±2 standard deviations. Figure 2B shows the same data, but expressed as a heat map of time-voltage values. Such heat maps can aid in the identification of multi-units (i.e., spikes from two or more neurons inadvertently lumped together in the same unit; see the script MLIB_Function_Reference.m for an example). Another indication for a multi-unit is a bimodal distribution of waveform peaks such as those shown in Figure 2C. The blue distribution represents the noise distribution for reference, the green (red) distribution represents all waveforms’ minimum (maximum) values. As this unit’s spike amplitude detection threshold was negative (vertical dotted line), primarily the green distribution is of relevance (for spikes with positive peaks, the red distribution would be more relevant). A Gaussian curve (black) is fitted to the amplitude distribution and serves to estimate the frequency of false negative spikes. Assuming normally distributed waveform amplitudes, there is a conspicuous lack of spikes in the rightmost tail of the distribution (arrow). For this example, it is estimated that 2% of spikes escaped detection. Figure 2D shows the inter-spike interval (ISI) distribution for this unit. ISIs of less than 3-4 ms indicate violation of the action potential refractory period and are not consistent with all spikes originating from the same neuron. For this unit, less than 0.5 % of spikes violate this criterion, which is a commonly used value to denote a unit as “single unit” (e.g., (Reyes-Puerta et al., 2015).

**Figure 2.**
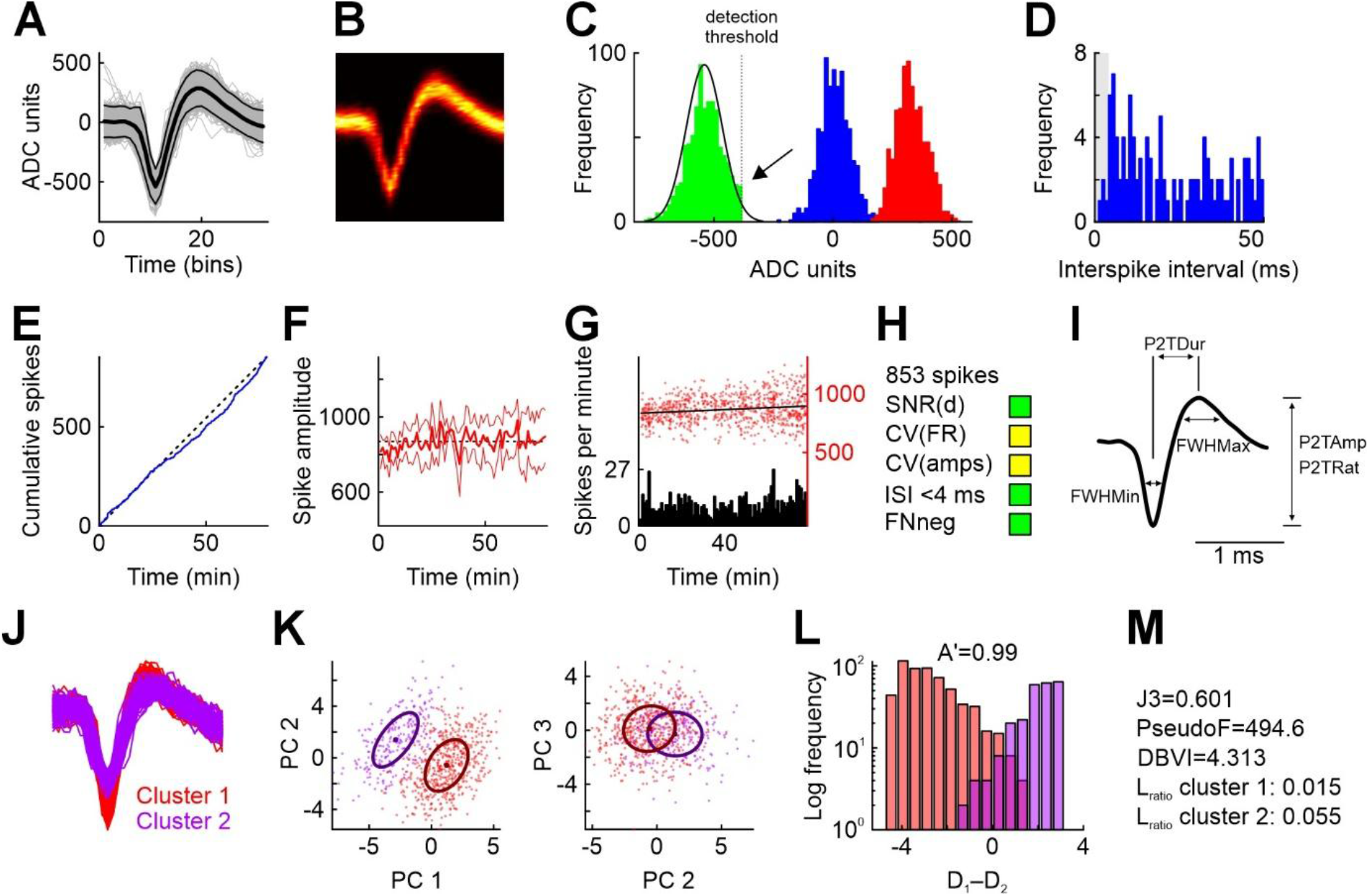
Illustration of some MLIB functions to examine spike sorting quality (functions mcc, mcheck, mwave). A) All 853 waveforms from a sparsely firing single unit recorded in the pigeon nidopallium caudolaterale (gray), mean waveform (thick black line) and ±2 standard deviations (thin black lines). ADC, analog-to-digital conversion (depends on the units of voltage supplied by the user. B) Time-voltage heat map of the waveforms in A. C) Distributions of baseline (blue) and peak voltages (green: minima, red: maxima). The black normal curve was fitted to the green distribution and is used to estimate false negatives. Due to a conservative detection threshold (dotted line), the green distribution appears truncated (arrow), leading to missed spikes (estimate false-negative rate: 2%). D) Inter-spike interval histogram. Gray shaded area highlights period of 4 ms in which no spikes are expected. The fraction of spikes occurring in this interval should be near zero for clean single units. E) Stability of firing can be assessed by plotting the cumulative spike count over session time. Diagonal black dotted line for comparison to expected steady firing rate. F) Stability of the unit can further be assessed by plotting the spike waveform amplitude averaged over 1-minute bins as a function of time (thick red line). Thin red lines represent ±1 standard deviation. G) Similar to E and F, but plotting binned spike counts (bin size 1 minute) and all spike amplitudes (red dots) as a function of session time. The regression line suggests an average increase of spike amplitude (which however amounts to only 0.8 ADC units per minute). H) Some quality indices returned by mcheck. SNR(d), signal-to-noise-ratio, measured as the distance between the maximum and the minimum of the average waveform, divided by the pooled 99%-Winsorized standard deviation. CV(FR) and CV(amps) represent the coefficient of variation of the firing rate and the amplitudes and serve as stability indicators (see documentation for details). I) Waveform features extracted with mwave. FWHMax and FWHMin, full width at half maximum and minimum, respectively. P2TDur, P2TRat, P2TAmp, peak-to-trough duration, ratio, and amplitude, respectively. J) Waveforms from two clusters of putative single units obtained from spike sorting. K) Waveforms plotted in principal component (PC) space. The first three PCs explain 44.8% of the total variance. Each dot represents a waveform. Bold points represent cluster means, ellipses represent the estimated covariance matrix (i.e., one standard deviation of a 2D Gaussian curve expressed as Mahalanobis distance in 2D space). L) Histogram of differences between the distance to the two cluster means for each waveform, plotted separately for the two clusters. Perfect cluster separation is achieved if the two distributions show zero overlap (i.e., no positive values of cluster 1 and no negative values of cluster 2). Class 1 and class 2 hit rates are proportions of correctly classified spikes for all possible values of D1–D2. Values of D can be understood as standard deviations in PC space, expressed as Mahalanobis distances. A’ represents the area under the ROC curve which can be maximally 1; excellent cluster separability is achieved at values of A’>0.98, while values <0.8 indicate very poor cluster separability. M) mcc returns several standard unit isolation quality measures. Panels A through I are modified from mcheck output, panels J through M from mcc output.

Figures 2E through 2G are useful to assess the stability of the recording. While action potential recordings with chronically implanted microwires are usually stable over several hours, acute recordings with single electrodes or electrode arrays tend to be less stable because of electrode drift or movements of the animal e.g. in head-fixed settings. Figure 2E shows cumulated spikes as a function of time; a perfectly stable unit would be expected to have a curve very close to the main diagonal of the figure (black dotted line). However, neuronal firing rates in many brain areas are expected to vary as a function of experimental parameters, e.g. when recording from behaving animals. In that case, cumulated spikes are less informative about recording stability, but the change in waveform amplitude is (shown in Figure 2F as a function of time). Figure 2G allows combined assessment of firing rate (black histogram with 1-min bins) and spike amplitude (red dots). mcheck also provides two quantitative measures to gauge unit stability (CV(amps) and CV(fr), which are explained in the function reference). Assessing unit stability is of paramount importance especially when neurons are observed under various conditions which change blockwise, as for example in some behavioral studies (Elber-Dorozko & Loewenstein, 2018). The function also provides some metrics useful for quantitative evaluation and selection of units for further analyses, along with three-stage recommendations (symbolized with traffic lights and easily customizable; Figure 2H). Finally, Figure 2I shows how the function mwave extracts basic waveform properties to assist in the classification into e.g. putative excitatory and inhibitory neurons (full width at half maximum, peak-to-trough duration etc.; (McCormick et al., 1985)).

While mcheck focuses on signal quality of a single waveform cluster, the function mcc allows to assess the separability of two or more spike clusters. In the present example, waveforms from a single electrode were clustered into two groups. Figure 2J shows spikes from the two clusters superimposed and color-coded for visual inspection. Waveforms from both clusters were subjected to principal component analysis, and Figure 2K shows the first three principal principal components for all waveforms. mcc offers several quantitative cluster separability indices. Figure 2L provides an intuitive depiction of the classification performance (along with the non-parametric area under the curve A’ which should approximate 1; a value of 0.5 signifies complete overlap of the two clusters). The x-axis represents the difference of the distances of each spike from its cluster center and to the center of the other cluster. If all spikes are closer to their respective cluster center than to that of the other cluster, the two distributions are completely separated and do not include 0, and A’=1 (note the logarithmic scaling of the ordinate). Other widely used quantitative indices provided by mcc include J3, PseudoF, DBVI, Lratio, and isolation distance (Nicolelis et al., 2003; Schmitzer-Torbert et al., 2005), see Figure 2M.

### Single-neuron spike train analysis

After spike sorting is finished, the next analysis step is typically to visualize the response characteristics of a unit by means of a peri-stimulus time histogram (PSTH) and an event-triggered raster display (Figure 3A, constructed with mpsth). As PSTHs are inherently noisy due to limited recording durations, the instantaneous firing rate is frequently estimated through convolution of the PSTH with a filtering kernel, yielding the so-called spike density function (SDF). Figure 3A shows an SDF obtained from convolution of the PSTH with an exponentially modified Gaussian kernel (yellow line) obtained through msdf (the function also offers boxcar, Gaussian, and simple exponential kernels of arbitrary width). Figure 3A also illustrates the output of the function mlat which can be used to estimate a unit’s response latency relative to a certain event (colored dots plotted over the peak of the SDF). mlat offers five common measures of response latency, and three of these are shown in Figure 3A.

**Figure 3.**
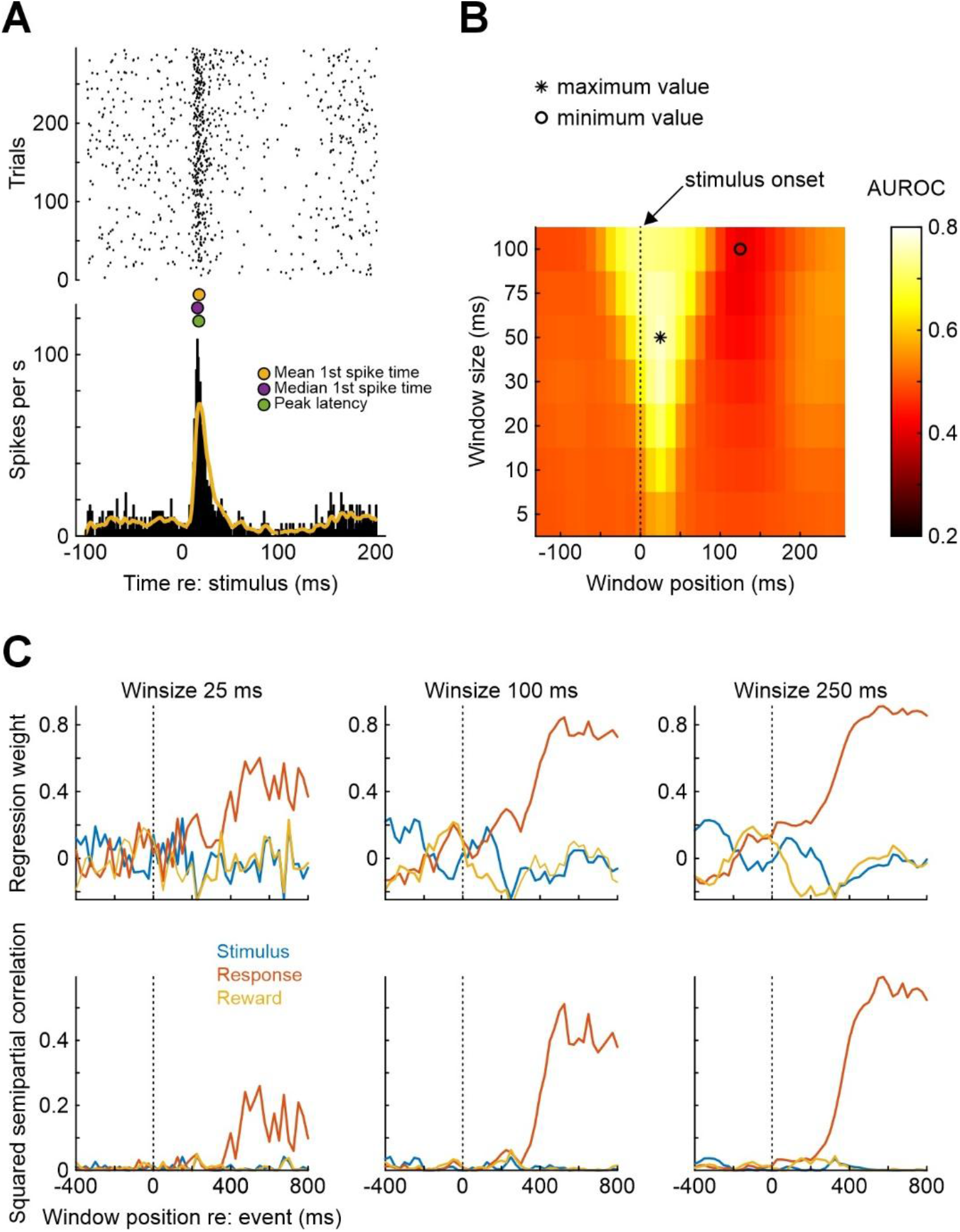
Illustration of some MLIB functions pertaining to single-neuron spike train analysis (functions mpsth, msdf, mlat, mwa, mmultregress). A) Peri-stimulus time histogram and raster plot of a single neuron from rat auditory cortex (mpsth). The yellow line represents a spike density function generated by msdf, using an exponentially modified Gaussian kernel with SD = 1 ms and tau = 5 ms. The three colored points between the two subpanels mark different estimates of response latency returned by mlat. B) Sliding window analysis results (mwa). C) Multiple regression analysis of spike counts on three different predictors (mmultregress). Spike counts of a rat medial frontal cortex neuron recorded during execution of a single-interval forced choice auditory discrimination task were subjected to regression analysis with the binary predictors stimulus, response, and reward, computed in successive windows shifted by 25 ms, with three different analysis window durations (25 ms, 100 ms, 250 ms). Spike counts were normalized before regression was performed. Top row shows the regression coefficients (intercept omitted), bottom row shows the squared semipartial correlation coefficient between each predictor and spike count, for each analysis window.

Especially in sensory neurophysiology, precise estimation of the timing and content of information in a spike train is of interest. This can be achieved through the functions mwa and mwa2. Figure 3B shows the output of mwa, computed for the same neuron as depicted in Figure 3A. Analysis windows of varying duration (ranging from 5 to 100 ms) were shifted in steps of 10 ms relative to the onset of the stimulus at time point 0. In each analysis window, the area under the receiver operating characteristic (ROC) curve (AUROC) is computed for spike counts in the analysis windows against spike counts in a reference window (here, pre-stimulus baseline) of the same duration. The heat map in Figure 3B shows that the maximum change in firing rate is present within 20-30 ms after stimulus onset. From a readout perspective, the optimal (AUROC-maximizing) analysis window duration is 50 ms, centered at +20 ms (AUROC=0.77; the window is highlighted by an asterisk in the figure). Such information is useful to understand how downstream brain areas might read out information from sensory representations for decision-making (Stüttgen & Schwarz, 2008, 2010). An alternative measure for AUROC is mutual information, which can be computed via the function mmi (Panzeri & Treves, 1996; Treves & Panzeri, 1995).

A frequently employed technique to examine the coding properties of single neurons is multiple regression. The general idea is that a neuron’s spike counts can be influenced by several factors which can be difficult to disentangle experimentally (e.g., (Kim et al., 2009; Romo et al., 2002). These factors can be entered as predictors, with spike counts as criterion variable, into a generalized linear model (GLM). For example, we recorded single neurons from rat dorsomedial frontal cortex (dMFC) while the animal was performing a two-stimulus auditory forced-choice discrimination task (Stoilova et al., 2020). In each of 160 trials, one out of two 100-ms auditory stimuli (noise bursts) differing in frequency content was presented when the animal’s snout entered a centrally located nose port. The animal had to wait until 50 ms after stimulus offset and then was free to withdraw and move into either of two laterally located choice ports. Following stimulus (S) 1, the animal received reward when entering the right choice port, following S2, reward was obtained when the animal entered the left choice port. The rats classified the stimuli correctly in around 80% of trials. We asked whether dMFC firing rates reflected stimulus identity, the choice response, or their interaction (reward or time-out punishment). Moreover, we asked at what time point(s) during the trial epochs dMFC neurons represent this information. We used the function mmultregress to perform this analysis, regressing single-neuron spike counts (the criterion variable) in 25-, 100-, and 250-ms windows shifted smoothly relative to stimulus onset (Figure 3C)^1^. Predictor variables were stimulus identity (S1 or S2), the choice response (right or left), and reward (yes or no; signed-effects coding, i.e., -1 vs. +1). The results are shown in Figure 3C. The top panels show the weights for the three regressors as a function of analysis window position, the bottom panels show the squared semipartial correlation coefficients (r_sp_^2^) of each predictor. It is evident that this neuron mainly represents the animal’s choice response (red curves), starting around 250 ms after stimulus onset. Regression weights and r_sp_^2^ values are generally larger and less noisy with longer analysis windows (at the expense of temporal resolution). Generally speaking, r_sp_ ^2^ represents the unique fraction of variance explained by any regressor, purified from its correlation with the other regressors, which can aid the interpretation of the effect of the independent variables (Cohen et al., 2003).

### Population analysis through dimensionality reduction

In recent years, due to technological advancements which allow to simultaneously record from large numbers of neurons (Steinmetz et al., 2021), the focus of neurophysiologists has shifted from the classic analysis of single-neuron spike trains and across-neuron averages to the visualization of population activity trajectories in multiple dimensions (Ebitz & Hayden, 2021; Saxena & Cunningham, 2019). The function mdimreduct accepts averaged spike counts from N neurons, performs PCA-based dimensionality reduction, and plots the resultant population activity trajectory in 2- and 3-dimensional space. Figure 4A visualizes the input array to the function – z-scored firing rates in 100-ms analysis windows for 300 dMFC neurons, aligned to the onset of stimulus 1 (same auditory discrimination task as described in the previous section). Neural activity was taken from 400 ms before until 400 ms after stimulus onset (correct trials only). This matrix was subjected to principal component analysis. Figure 4B shows the scores of the first 16 principal components (PCs), Figure 4C shows explained variance cumulated over the first 16 PCs. As frequently observed with this kind of data, the 300-dimensional activity vector which served as the input can be safely reduced to a handful of PCs (the first 5 PCs explain almost 80% of the total variance). Figure 4D shows the population trajectory for the first three PCs for correct S1 trials. Time relative to stimulus onset is color-coded, with darker points representing earlier and lighter dots representing later time points. When the data matrix from Figure 4A for S1 is concatenated with the corresponding matrix for S2 (not shown) and the concatenated matrix is then subjected to PCA, population trajectories for the two types of trials – S1 presentations followed by movement to the right, S2 presentations followed by movement to the left – can be visualized together as in Figure 4E. Before stimulus onset, the two trajectories are virtually identical. Shortly after stimulus onset, the two trajectories diverge (arrow) as the animal makes its decision to move to either of the two choice ports. This divergence is observed already before the stimulus has ended and the animal has withdrawn from the center port, indicating early involvement of the dMFC population in the decision process and/or response planning.

**Figure 4.**
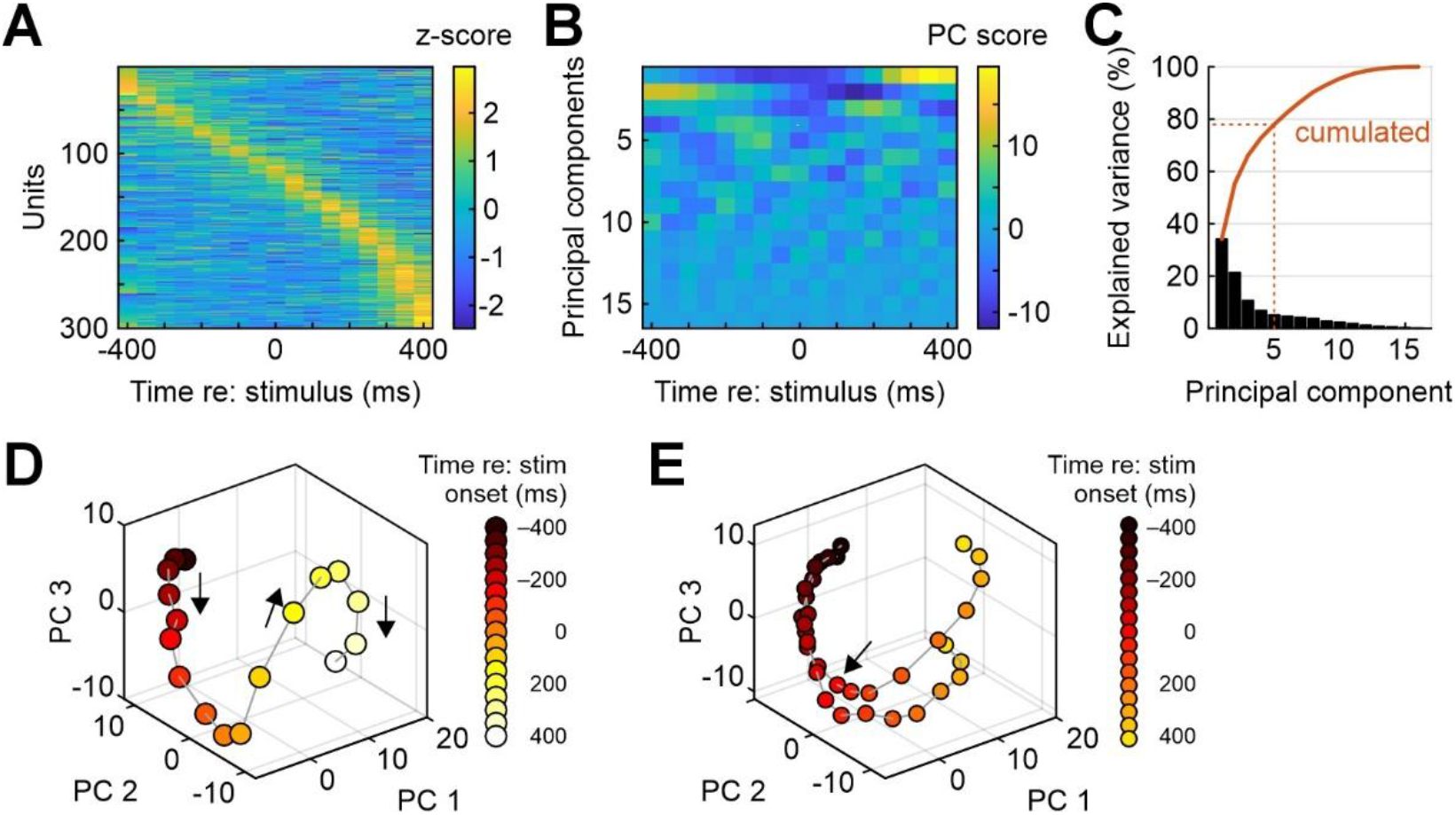
Illustration of dimensionality reduction for multi-neuron recordings (function mdimreduct). A) Average neuronal activity (z-scored and color-coded) for 300 neurons from medial frontal cortex around onset of an auditory stimulus (one of two), sorted by peak latency. B) The first 16 principal components (PCs) computed from the data in panel A. C) Percentage explained variance by the first 16 principal components (bars). Red line shows cumulated percentage. The first five PCs together explain roughly 78% of the variance. D) Population trajectory for the first three PCs. Time is color-coded from black (-400 ms relative to stimulus onset) to white (+400 ms). E) Same as panel D, but after dimensionality reduction for both stimuli together. The trajectories for the two types of trials begin to diverge shortly after stimulus onset (time point 0 marked by arrow).

### Analysis of local field potentials

While the majority of MLIB functions are concerned for spike train analysis, three functions are useful for analyzing local field potential (LFP) data (mcsd, mep, mspectan). Here, I will introduce only the function mcsd which computes and plots the results of a current-source density (CSD) analysis. CSD is a useful tool to assign individual contacts from linear electrode arrays to cortical layers (Vandevelde et al., 2023; Yang et al., 2017; Yeganeh et al., 2022). Figure 5A depicts averaged LFP (voltage) traces from 20 linearly spaced electrodes (inter-electrode distance 50 µm) inserted into the barrel cortex of an anesthetized mouse (van der Bourg et al., 2017). The position of the electrode is color-coded (black: topmost electrode contact, white: lowermost electrode contact). A single sinusoidal whisker stimulus was applied, starting at time point 0. LFPs were averaged over 20 trials. Figure 5B shows the same data as in 4A as a heat map, highlighting that the strongest LFP response was observed at the 5th contact from the top. Figure 5C shows the raw CSD (i.e., the second spatial derivative of the LFP in Figure 5B). The raw CSD was then spatially interpolated 10-fold and smoothed (Figure 5D) to allow easier identification of the earliest current sink (white arrow) which can be used to identify layer IV, allowing precise assignment of electrode contacts (and the units recorded with them). The other electrode contacts can then be assigned to the remaining layers through the combined consideration of the current sources and histological references (Mitzdorf, 1985; Reyes-Puerta et al., 2015).

**Figure 5.**
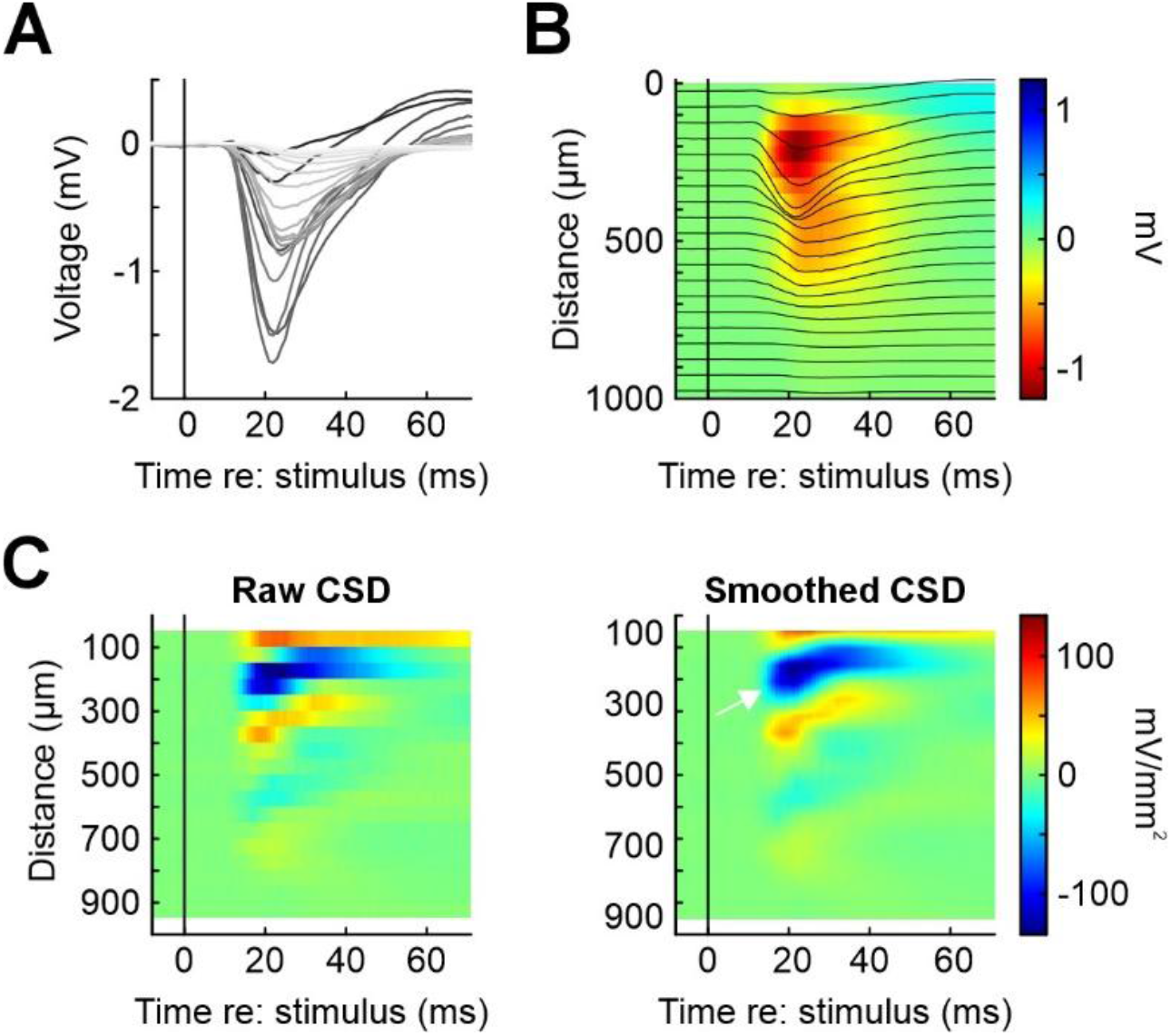
Illustration of MLIB function mcsd for LFP data. A) Laminar LFP recordings in mouse barrel cortex after deflection of the principal whisker. Each line represents the average over 10 stimulus repetitions for one of 20 electrodes from a linear array (electrode spacing 50 µm). B) Same data as in panel A, but plotting distance of the electrodes from the pial surface on the ordinate. Average LFP responses are shown both as lines and pseudocolored (in mV). C) Current source density (CSD) plots for the data in panels A and B. Left: raw CSD, right: CSD after 10-fold spatial interpolation. Pseudocolor scale units are mV/mm2. This data was collected by Dr Jenq-Wei Yang in the lab of Prof. Heiko Luhmann, University Medical Center Mainz, and is part of a larger data set published in (van der Bourg et al., 2017). The authors kindly granted permission to use this data in the present publication.

## Discussion

In summary, MLIB is an easy-to-use yet versatile open-source toolbox for the analysis of extracellular spike and field potential data. Through detailed documentation and usage demonstrations in an accompanying script (MLIB_Function_Reference.m), it should be useful for both beginners and more experienced experimentalists in need of basic as well as some more advanced analysis functions. For more specialized analysis techniques, especially for large-scale neuronal recordings, the reader is referred to (Stringer et al., 2025; Stringer & Pachitariu, 2024); but note that their codes are written in Python. For comprehensive overviews of spike and LFP analysis toolboxes written in Matlab and Python, see (Unakafova & Gail, 2019) and the SpikeResources collection on GibHub (https://github.com/openlists/SpikeResources).

There exist many other toolboxes for electrophysiological data analysis, mainly within the Matlab and Python environments, all of them sharing some overlap while serving somewhat different purposes. MLIB has several features which set it apart from other neurophysiology toolboxes. First, I believe that a particular strength of MLIB is that it offers a broad and integrated workflow, covering a wide array of standard analysis steps that range from spike sorting quality assessment to the visualization of activity to more specialized analyses such as information content to dimensionality reduction. This workflow is illustrated in Figure 1 for didactic purposes. MLIB therefore is comprehensive in the sense that many labs may need no or only few additional tools for a thorough analysis of their electrophysiological data to meet current standards in the field. This distinguishes MLIB from more specialized toolboxes such as (Python-based) SpikeInterface (Buccino et al., 2020) which focuses on spike sorting, or Chronux (Bokil et al., 2010), which is based on Matlab and is not only highly useful for spike sorting but also provides for advanced spectral analysis of LFP, spike trains, and their coherence. In the Matlab programming environment, other useful tools are FieldTrip (Oostenveld et al., 2011), which is a comprehensive, continuously curated and expanded freely available Matlab-based software package for MEG, EEG, and iEEG analysis. However, both of these packages do not offer many functions for spike train visualization and analysis. For users wanting to avoid coding altogether, CellExplorer (Petersen et al., 2021) which offers a comprehensive, interactive analysis environment with a graphical user interface, standardized data structures for inter-laboratory data pooling and comparison, focused on single-neuron characterization, especially in terms of defined cell types and their biophysical properties.

Second, MLIB by design specifically addresses newcomers to neurophysiology with minimal programming skills. The functions are simple one-liners and accompanied by extensive documentation and examples in a reference file. The basic idea is to have one analysis in one function. In consequence, the MLIB functions can easily be incorporated within analysis scripts – their output can for example be collected for several units are datasets and then submitted to further, custom analyses tailored to the specific needs of the experimenter. This contrasts with toolboxes such as Elephant (Denker et al., 2018) or the FMAToolbox within Neurosuite (https://neurosuite.sourceforge.net/) which impose somewhat higher demands on programming knowledge and sometimes return results in more complicated data structures, while MLIB outputs are, for the most part, scalars, vectors, and matrices, and thus easy to handle.

There are several limitations of this toolbox. First and foremost, it is written in Matlab, which is a proprietary language and therefore requires a running Matlab license. Additionally, many of the functions require active licenses for one or more toolboxes, which can be costly. This contrasts with open-source alternatives such as Elephant (Denker et al., 2018)or CellExplorer (Petersen et al., 2021) which rely on Python. On the other hand, Matlab offers several assets, such as user-friendliness, an excellent documentation, as well as an active community on Matlab Central File Exchange (https://de.mathworks.com/matlabcentral/fileexchange). Also, cursory inspection of electrophysiology papers shows that many labs are using Matlab and are likely to continue to do so for the foreseeable future. The second limitation of MLIB is that it does not contain spike-sorting functions. That said, there exists already a large variety of both Matlab- and Python-based spike sorting packages (e.g., (Pachitariu et al., 2024; Quian Quiroga et al., 2009; Rossant et al., 2016; Strohl et al., 2021; Ul Hassan et al., 2021)), as well as commercial alternatives (such as Plexon’s Offline Sorter). Anyway, each of these package’s outputs will eventually be spike-sorted waveforms and spike timestamps, which can then readily be used with MLIB. For example, our lab is using Offline Sorter and Spike2 for spike sorting, and we extensively check and validate sorting results with mcheck and mcc (e.g., Stoilova et al., 2020; Vandevelde et al., 2023). The third limitation is that MLIB is not geared towards large channel counts such as offered by Neuropixels probes. Functions such as mcheck and mcc are intended for single-channel recordings, allowing close inspection of waveform characteristics and spike sorting quality for manual curation (see above), but when large numbers of units are recorded, checking each and every unit’s sorting quality manually will become cumbersome. That argument notwithstanding, our lab has used MLIB functions mcheck and mcc with data sets of up to 700 units (and there were no problems or unacceptable processing times employing standard desktop computers). However, for most of MLIB’s functions, the number of recording channels and/or recorded units do not matter – PSTHs and SDFs are necessarily computed for one unit at a time, and spike count regression as well as population trajectories can be obtained for an unlimited number of units in principle (in my lab, we have used mmultregress and mdimreduct for datasets containing up to 700 units).

With the advent of large-scale multi-neuron recording techniques, analysis techniques will continue to evolve (Stringer & Pachitariu, 2024). However, the basic visualization and analysis of the single-neuron response properties is likely to remain an important starting point for the analysis of electrophysiological data. I hope that experimentalists will continue to appreciate the streamlined, easy-to-use modular functions provided by MLIB to get started exploring their spike data.

## Acknowledgements

The author is grateful for the feedback and support of present and former lab members. The LFP data were provided by Dr Jenq-Wei Yang and collected in the lab of Prof Heiko Luhmann, Institute of Physiology, University Medical Center Mainz, and are used with the authors’ permission.

## Funding

Preparation of the toolbox and the manuscript were supported by several grants from the Deutsche Forschungsgemeinschaft to MCS (project IDs 179167628, 197059818, 223368015, 280347763, 424828846, and 543128862).

## Conflict of interest disclosure

The authors declare that they comply with the PCI rule of having no financial conflicts of interest in relation to the content of the article.

## Data, scripts, code, and supplementary information availability

Scripts, data, code, and additional information are available online: https://github.com/maikstue/mlib-spike-data

and

https://de.mathworks.com/matlabcentral/fileexchange/37339-mlib-toolbox-for-analyzing-spike-data.

and

https://doi.org/10.5281/zenodo.19455823

## Footnotes

1 In this example, regression was performed assuming a normally distributed response variable, but spike count distributions may be better described with Poisson distributions, especially if only few spikes are observed in each analysis time bin. Since the function mmultregress uses Matlab’s fitglm, it also accommodates the Poisson distribution (set input argument ‘udist’ to ‘poisson’). In the present case, the results are qualitatively similar.

## Notes

https://github.com/maikstue/mlib-spike-data

https://de.mathworks.com/matlabcentral/fileexchange/37339-mlib-toolbox-for-analyzing-spike-data

https://doi.org/10.5281/zenodo.19455823

### Competing Interest Statement

The authors have declared no competing interest.

### Summary of Updates

The previous version has meanwhile received a positive recommendation from PCI Neuroscience, see https://doi.org/10.24072/pci.neuro.100467 and will be published in Peer Community Journal.

## References

Adrian, E. D. (1926). The impulses produced by sensory nerve endings. Part I. Journal of Physiology, 61(1), 49–72.

Bokil, H., Andrews, P., Kulkarni, J. E., Mehta, S., & Mitra, P. P. (2010). Chronux: A platform for analyzing neural signals. Journal of Neuroscience Methods, 192(1), 146–151. 10.1016/j.jneumeth.2010.06.020

Buccino, A. P., Hurwitz, C. L., Garcia, S., Magland, J., Siegle, J. H., Hurwitz, R., & Hennig, M. H. (2020). SpikeInterface, a unified framework for spike sorting. ELife, 9, 1–24. 10.7554/eLife.61834

Chakrabarti, S. (2024). Electrophysiology Tutorial for Neuroscience. <https://github.com/MathWorks-Teaching-Resources/Electrophysiology-Tutorial-for-Neuroscience/releases/tag/v1.2 >

Cohen, J., Cohen, P., West, S. G., & Aiken, L. S. (2003). Applied multiple regression/correlation analysis for the behavioral sciences (3rd ed.). Taylor & Francis Group.

DeCharms, R. C., & Merzenich, M. M. (1996). Primary cortical representation of sounds by the coordination of action-potential timing. Nature, 381, 610–613.

Denker, M., Yegenoglu, A., & Grün, S. (2018). Collaborative HPC-enabled workflows on the HBP Collaboratory using the Elephant framework. In Neuroinformatics (p. 19).

Ebitz, R. B., & Hayden, B. Y. (2021). The population doctrine in cognitive neuroscience. Neuron, 109(19), 3055–3068. 10.1016/j.neuron.2021.07.011

Elber-Dorozko, L., & Loewenstein, Y. (2018). Striatal Action-Value Neurons Reconsidered. ELife, 7, e34248. 10.1101/087502

Hill, D. N., Mehta, S. B., & Kleinfeld, D. (2011). Quality metrics to accompany spike sorting of extracellular signals. Journal of Neuroscience, 31(24), 8699–8705. 10.1523/JNEUROSCI.0971-11.2011

Kasties, N., Starosta, S., Güntürkün, O., & Stüttgen, M. C. (2016). Neurons in the pigeon caudolateral nidopallium differentiate Pavlovian conditioned stimuli but not their associated reward value in a sign-tracking paradigm. Scientific Reports, 6(October), 35469. 10.1038/srep35469

Kim, H., Sul, J. H., Huh, N., Lee, D., & Jung, M. W. (2009). Role of striatum in updating values of chosen actions. Journal of Neuroscience, 29(47), 14701–14712. 10.1523/JNEUROSCI.2728-09.2009

Kleinfeld, D., Luan, L., Mitra, P. P., Robinson, J. T., Sarpeshkar, R., Shepard, K., Xie, C., & Harris, T. D. (2019). Can One Concurrently Record Electrical Spikes from Every Neuron in a Mammalian Brain? Neuron, 103(6), 1005–1015. 10.1016/j.neuron.2019.08.011

Lengersdorf, D., Pusch, R., Güntürkün, O., & Stüttgen, M. C. (2014). Neurons in the pigeon nidopallium caudolaterale signal the selection and execution of perceptual decisions. The European Journal of Neuroscience, 40(9), 3316–3327. 10.1111/ejn.12698

Lettvin, J. Y., Maturana, H. R., McCulloch, W. S., & Pitts, W. H. (1959). What the frog’s eye tells the frog’s brain. Proceedings of the IRE, 47(11), 1940–1951.

McCormick, D. A., Connors, B. W., Lighthall, J. W., & Prince, D. A. (1985). Comparative Electrophysiology of Pyramidal and Sparsely Spiny Stellate Neurons of the Neocortex. Journal of Neurophysiology, 54(4), 782–806.

Mitzdorf, U. (1985). Current source-density method and application in cat cerebral cortex: investigation of evoked potentials and EEG phenomena. Physiological Reviews, 65(1), 37–100. https://doi.org/n/a

Nicolelis, M. A. L., Dimitrov, D., Carmena, J. M., Crist, R., Lehew, G., Kralik, J. D., & Wise, S. P. (2003). Chronic, multisite, multielectrode recordings in macaque monkeys. Proceedings of the National Academy of Sciences of the United States of America, 100(19), 11041–11046. 10.1073/pnas.1934665100

Oostenveld, R., Fries, P., Maris, E., & Schoffelen, J. M. (2011). FieldTrip: Open source software for advanced analysis of MEG, EEG, and invasive electrophysiological data. Computational Intelligence and Neuroscience, 2011. 10.1155/2011/156869

Pachitariu, M., Sridhar, S., Pennington, J., & Stringer, C. (2024). Spike sorting with Kilosort4. Nature Methods, 21(5), 914–921. 10.1038/s41592-024-02232-7

Panzeri, S., & Treves, A. (1996). Analytical estimates of limited sampling biases in different information measures. Network: Computation in Neural Systems, 7(1), 87–107. 10.1080/0954898x.1996.11978656

Petersen, P. C., Siegle, J. H., Steinmetz, N. A., Mahallati, S., & Buzsáki, G. (2021). CellExplorer: A framework for visualizing and characterizing single neurons. Neuron, 109(22), 3594–3608.e2. 10.1016/j.neuron.2021.09.002

Quian Quiroga, R., Nadasdy, Z., & Ben-Shaul, Y. (2009). Unsupervised spike detection and sorting with wavelets and superparamagnetic clustering. Neural Computation, 16(8), 1661–1687. 10.1162/089976604774201631

Reyes-Puerta, V., Sun, J.-J., Kim, S., Kilb, W., & Luhmann, H. J. (2015). Laminar and columnar structure of sensory-evoked multineuronal spike sequences in adult rat barrel cortex in vivo. Cerebral Cortex, 25(8), 2001–2021. 10.1093/cercor/bhu007

Romo, R., Hernández, A., Zainos, A., Lemus, L., & Brody, C. D. (2002). Neuronal correlates of decision-making in secondary somatosensory cortex. Nature Neuroscience, 5(11), 1217–1225.

Rossant, C., Kadir, S. N., Goodman, D. F. M., Schulman, J., Hunter, M. L. D., Saleem, A. B., Grosmark, A., Belluscio, M., Denfield, G. H., Ecker, A. S., Tolias, A. S., Solomon, S., Buzski, G., Carandini, M., & Harris, K. D. (2016). Spike sorting for large, dense electrode arrays. Nature Neuroscience, 19(4), 634–641. 10.1038/nn.4268

Salinas, E., & Thier, P. (2000). Gain modulation: a major computational principle of the central nervous system. Neuron, 27(1), 15–21.

Saxena, S., & Cunningham, J. P. (2019). Towards the neural population doctrine. Current Opinion in Neurobiology, 55, 103–111.

Schmitzer-Torbert, N., Jackson, J., Henze, D., Harris, K., & Redish, A. D. (2005). Quantitative measures of cluster quality for use in extracellular recordings. Neuroscience, 131(1), 1–11. 10.1016/j.neuroscience.2004.09.066

Starosta, S., Stüttgen, M. C., & Güntürkün, O. (2014). Recording single neurons’ action potentials in freely moving pigeons across three stages of learning. Journal of Visualized Experiments, 88, e51283.

Steinmetz, N. A., Aydin, C., Lebedeva, A., Okun, M., Pachitariu, M., Bauza, M., Beau, M., Bhagat, J., Böhm, C., Broux, M., Chen, S., Colonell, J., Gardner, R. J., Karsh, B., Kostadinov, D., Mora-Lopez, C., Park, J., Putzeys, J., Sauerbrei, B., … Harris, T. D. (2021). Neuropixels 2.0: A miniaturized high-density probe for stable, long-term brain recordings. Science, 372. 10.1101/2020.10.27.358291

Stoilova, V. V, Knauer, B., Berg, S., Rieber, E., Jäkel, F., & Stüttgen, M. C. (2020). Auditory cortex reflects goal-directed movement but is not necessary for behavioral adaptation in sound-cued reward tracking. Journal of Neurophysiology, 124(4), 1056–1071. 10.1152/jn.00736.2019

Stringer, C., & Pachitariu, M. (2024). Analysis methods for large-scale neuronal recordings. Science, 386(6722). 10.1126/science.adp7429

Stringer, C., Zhong, L., Syeda, A., Du, F., Kesa, M., & Pachitariu, M. (2025). Rastermap: a discovery method for neural population recordings. Nature Neuroscience, 28(1), 201–212. 10.1038/s41593-024-01783-4

Strohl, J. J., Gallagher, J. T., Gómez, P. N., Glynn, J. M., & Huerta, P. T. (2021). Framework for automated sorting of neural spikes from Neuralynx-acquired tetrode recordings in freely-moving mice. Bioelectronic Medicine, 7(1), 17. 10.1186/s42234-021-00079-3

Stüttgen, M. C., & Schwarz, C. (2008). Psychophysical and neurometric detection performance under stimulus uncertainty. Nature Neuroscience, 11(9), 1091–1099.

Stüttgen, M. C., & Schwarz, C. (2010). Integration of vibrotactile signals for whisker-related perception in rats is governed by short time constants: comparison of neurometric and psychometric detection performance. Journal of Neuroscience, 30(6), 2060–2069.

Treves, A., & Panzeri, S. (1995). The Upward Bias in Measures of Information Derived from Limited Data Samples. Neural Computation, 7(2), 399–407. 10.1162/neco.1995.7.2.399

Ul Hassan, M., Veerabhadrappa, R., & Bhatti, A. (2021). Efficient neural spike sorting using data subdivision and unification. PLOS ONE, 16(2), e0245589. 10.1371/journal.pone.0245589

Unakafova, V. A., & Gail, A. (2019). Comparing Open-Source Toolboxes for Processing and Analysis of Spike and Local Field Potentials Data. In Frontiers in Neuroinformatics (Vol. 13). Frontiers Media S.A. 10.3389/fninf.2019.00057

van der Bourg, A., Yang, J.-W., Reyes-Puerta, V., Laurenczy, B., Wieckhorst, M., Stüttgen, M. C., Luhmann, H. J., & Helmchen, F. (2017). Layer-Specific Refinement of Sensory Coding in Developing Mouse Barrel Cortex. Cerebral Cortex, 27, 4835–4850. 10.1093/cercor/bhw280

Vandevelde, J. R., Yang, J.-W., Albrecht, S., Lam, H., Kaufmann, P., Luhmann, H. J., & Stüttgen, M. C. (2023). Layer- and cell-type-specific differences in neural activity in mouse barrel cortex during a whisker detection task. Cerebral Cortex, 33, 1361–1382. 10.1093/cercor/bhac141

Yang, J.-W., Prouvot, P.-H., Reyes-Puerta, V., Stüttgen, M. C., Stroh, A., & Luhmann, H. J. (2017). Optogenetic Modulation of a Minor Fraction of Parvalbumin-Positive Interneurons Specifically Affects Spatiotemporal Dynamics of Spontaneous and Sensory-Evoked Activity in Mouse Somatosensory Cortex in Vivo. Cerebral Cortex, 27(12), 5784–5803. 10.1093/cercor/bhx261

Yeganeh, F., Knauer, B., Backhaus, R. G., Yang, J. W., Stroh, A., Luhmann, H. J., & Stüttgen, M. C. (2022). Effects of optogenetic inhibition of a small fraction of parvalbumin - positive interneurons on the representation of sensory stimuli in mouse barrel cortex. Scientific Reports, 12, 19419. 10.1038/s41598-022-24156-y

